# Frame-specific depletion of the TRBV23-1 pseudogene in human TCR repertoires: Quantitative evidence and possible biological explanations

**DOI:** 10.1101/2025.09.30.679533

**Authors:** Jason Tigg, Anree Bektashi-Brown

## Abstract

V(D)J recombination generates T cell receptor diversity, but most rearrangements introduce frameshifts or premature termination codons that prevent formation of functional receptors. A quantitative model was developed to estimate the expected ratio of in-frame to out-of-frame arrangements. A statistical model was used to identify repertoires with anomalous frame usage. Applied to more than 6,000 repertoires from nine cohorts, the statistical model identified 14 repertoires with frame-dependent depletion in rearrangements involving the pseudogene TRBV23-1. The depletion is inconsistent with sequencing artifacts, contamination, or conventional thymic selection, but may reflect a biological process such as HLA class I-mediated targeting of TRBV23-1-derived peptides. RNA-seq analysis shows frame-specific RNA levels correlated with depletion rates, supporting the possibility of a post-transcriptional regulatory contribution. These observations suggest that pseudogenes, long considered inert, may influence immune repertoire structure. The findings establish a robust statistical signal of non-random TRBV23-1 depletion and provide testable mechanistic hypotheses for future validation.

## Introduction

Adaptive immune diversity is generated through V(D)J recombination, where V, D, and J gene segments are somatically rearranged with additional nucleotide trimming and insertions to create vast sequence variability (1–3). In T cells, this process produces the *α* and *β* chains of the *αβ* T cell receptor (TCR). Most rearrangement attempts fail to produce a functional receptor due to frameshifts or premature termination codons (PTCs) (4–6), in which case a second rearrangement attempt is probabilistically possible on the homologous chromosome (7).

Post V(D)J recombination, thymic selection eliminates thymocytes that do not recognize self-MHC or are overly self-reactive (8, 9). While productive rearrangements drive immune function, non-productive sequences provide an unbiased record of recombination dynamics and are valuable for modelling generative probabilities, allele usage, and repertoire structure (5, 10).

Traditionally, non-productive sequences arising from unsuccessful V(D)J recombination attempts have been considered inert by-products—genomic ‘baggage’ accompanying a successful rearrangement on the homologous chromosome. Their transcripts are rapidly degraded through mechanisms such as nonsense-mediated decay (NMD) (11), and thus they are generally dismissed as biologically irrelevant. However, emerging evidence suggests that pseudogenes and non-coding genes may retain regulatory influence, for example through transcript stability, peptide presentation, or other indirect pathways (12–14). Importantly, such functions have only been described for pseudogenes in general, and there is currently no evidence that TRBV pseudogenes themselves exert comparable regulatory roles. Whether such non-productive rearrangements, particularly those involving pseudogenes, might nevertheless exert influence on the immune system remains an open question. Addressing this question requires large-scale repertoire data spanning diverse biological contexts. In this work, such data are drawn from ten publicly available TCR sequencing cohorts, comprising more than 6,000 repertoires (see Table 5, Methods).

## Results

In this study, each unique CDR3 sequence in a repertoire is counted once, irrespective of its level of clonal expansion. Expansion reflects antigen-driven proliferation of the paired productive receptor, whereas the focus here is on the accompanying non-productive rearrangements.

### Notation

During translation, nucleotides are read in triplets (codons), and the starting position defines the reading frame. A rearrangement is labelled ‘in-frame’ when the total CDR3 length after trimming and insertions is a multiple of three nucleotides, preserving codon alignment. Other rearrangements are labelled ‘out-of-frame’. For example, 42 nucleotides is in-frame, whereas 43 or 44 correspond to the two out-of-frame cases. If the CDR3 length is not a multiple of three, the reading frame shifts, altering all downstream codons. Biologically, there is no clear rationale to distinguish between the two out-of-frame categories (multiples of three plus one versus multiples of three plus two). However, in this study all three frames are analyzed separately, enabling detection of frame-specific patterns that would otherwise be overlooked. To formalize the analysis, a clear notation is introduced to represent frame categories clearly and unambiguously.

Frames are denoted by *F*_0_, *F*_1_, and *F*_2_, where *F*_0_ represents the in-frame case (CDR3 nucleotide length divisible by three) and *F*_1_ and *F*_2_ represent the two out-of-frame cases. **Formally, the subscript is defined as the CDR3 nucleotide length modulo 3**. *F*_1_ and *F*_2_ are always non-productive. A necessary but not sufficient condition for *F*_0_ to be productive is that the CDR3 generative process did not introduce a termination codon. Even then, functionality depends on the chosen V-gene: genes incapable of generating a functional TCR, even with an *F*_0_ no-termination codon CDR3, are designated **pseudogenes**.

It is sometimes useful to split *F*_0_ into two subsets:

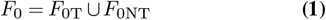

where *F*_0T_ denotes in-frame sequences with a PTC and *F*_0NT_ denotes those without.

For shorthand, combinations of frames can be written as unions using a comma-separated notation, e.g.

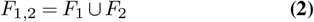

Finally, the union of all frames is denoted by

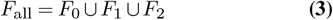

### Modelling the in-frame/out-of-frame ratio

As stated above, this study uniquely considers differences between *F*_0_, *F*_1_ and *F*_2_. Before turning to those distinctions, it is useful to begin with a more conventional analysis, in which the two out-of-frame rearrangements are grouped together. Modelling and empirical measurement begin with the following ratio, considered on a V-gene–specific basis:

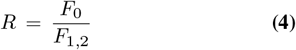

A general framework for V(D)J recombination is developed in the Methods section, incorporating parameters for thymic selection, gene usage probabilities, and other sources of biological variation. From this model a formula for the ratio *R* is obtained. Under simplifying assumptions, for functional genes the formula reduces to a tractable form with only three parameters: *p*_2_, the probability of a second recombination attempt after an unsuccessful first; *f*, the fraction of V genes that are pseudogenes; and *p*_T_, the probability that an in-frame rearrangement contains a PTC in the CDR3 region.

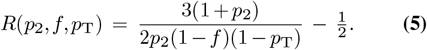

Using empirical estimates for humans (*f ≈* 0.33, *p*_T_ *≈* 0.18), the predicted ratios are given in Table 1 for various values of *p*_2_. With realistic parameters, functional V genes are expected to exhibit in-frame to out-of-frame ratios in the range of 5–11. By contrast, pseudogenes remain fixed at *R* = 0.5 in the absence of frameshift–dependent thymic selection pressures.

**Table 1.** Predicted in-frame/out-of-frame ratios (*R*) for functional genes and various probabilities *p*_2_ of a second recombination attempt.

| $p_2$ | $R$ |
| --- | --- |
| 0.3 | 11.3 |
| 0.6 | 6.8 |
| 0.8 | 5.7 |
| 1.0 | 5.0 |

To compare the theoretical prediction with empirical data, aggregated in-frame and out-of-frame sequence counts were computed for each V gene over the repertoires in Cohort C1 (see Methods for details). These counts are plotted in Figure 1 together with the pseudogene line (*y* = 0.5*x*) and an illustrative functional line (*y* = 6*x*). The slope of the functional line lies within the predicted range of 5–11 from Table 1, demonstrating qualitative agreement between theory and data.

**Fig. 1.**
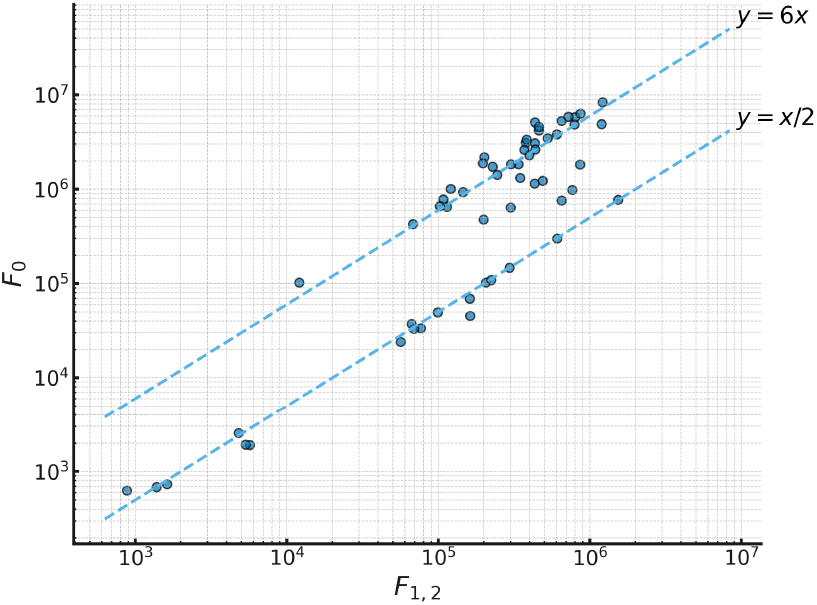
In-frame versus out-of-frame unique sequence counts by V gene across all repertoires in cohort C1. The pseudogene line (*y* = 0.5*x*) and a functional reference line (*y* = 6*x*) are shown for comparison.

Figure 2 shows a close-up that ignores V-genes with low sequence counts (fewer than 20,000 distinct sequences for *F*_1,2_ over the entire cohort). The top three lines on this plot correspond to different probabilities of a second recombination attempt (*p*_2_ values of 0.3, 0.6 and 0.8 respectively), providing direct comparison between model predictions and empirical data. The bottom line shows the pseudogene relation *y* = 0.5*x*.

**Fig. 2.**
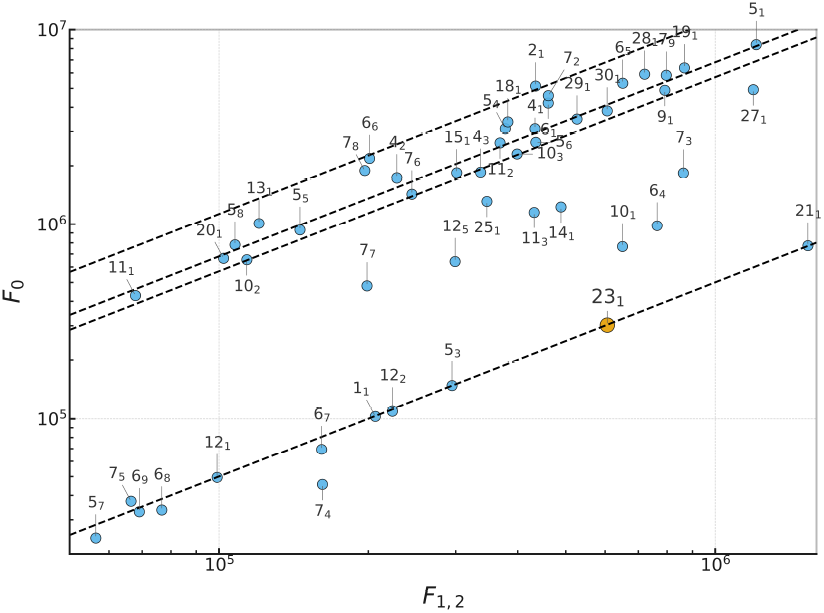
Enlarged view of Figure 1, highlighting the majority of V genes. Also included is the pseudogene line and three functional-gene lines, corresponding to *p*_2_ values of 0.3, 0.6, and 0.8 (from top to bottom).

At this point it is useful to pause and examine a couple of specific V genes in detail, to introduce a visualization that will be used extensively throughout the paper. For each gene a histogram of unique sequence counts by CDR3 length in nucleotides is shown, accompanied by a small inset displaying the overall frame counts (*F*_0_, *F*_1_, and *F*_2_). In the main histogram and the inset, the *F*_0_ bar is divided into *F*_0*T*_ (top) and *F*_0*NT*_ (bottom). While the inset summarises the overall balance between frames, the full histogram provides crucial additional information.

Technical artifacts, for example, have a distinctive signature which is clearly visible in the main histogram; specifically, a sharp spike of unique counts at a single CDR3 length (Figure 3). If only frame totals were considered, this might be misinterpreted as enhanced frequency of sequences in the affected frameshift or reduced frequency in the complementary frameshifts. Because sequencing reads often capture only part of the V-gene, long CDR3 sequences can result in ambiguous V-gene assignment, producing asymmetric or truncated distributions. More generally, any imbalance in frame usage that is not distributed across CDR3 lengths according to the canonical distribution signals the need for closer investigation. When unique sequence counts are plotted by CDR3 length, periodic patterns are immediately visible, offering an intuitive sense of statistical significance that a summary ratio or p-value alone may struggle to convey.

**Fig. 3.**
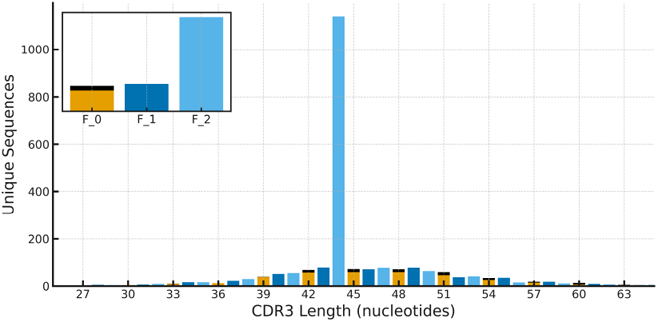
Unique TRBV23-1 sequence counts by CDR3 length for sample 03855000012652 in cohort C2. The massive spike observed in *F*_2_ arises entirely from sequences of a uniform CDR3 length (44 in this example). These sequences display hallmarks of technical artifacts: extreme GC content with long polyG tracts, near-identical nucleotides, a shared fixed suffix, and absence of canonical V/J motifs. Such features are characteristic of primer/adapter concatemers, vector or polylinker carryover, or synthetic oligonucleotides spuriously aligning to partial V/J regions, and can therefore be systematically identified and excluded from downstream analyses.

Full distribution histograms in combination with inset summaries can be regarded as both a quality-control tool and a revealing lens on frame-dependent biology. By way of example, two specific V genes are examined. Figure 4 displays the profile of the functional gene TRBV5-1, exhibiting the expected pattern of a strong peak in *F*_0_ and much smaller, commensurate peaks in *F*_1_ and *F*_2_. Figure 5 presents the pseudogene TRBV23-1, which is the focus of this paper. In this case the distribution is smooth, with equal representation across all three frameshifts, as expected for a pseudogene.

**Fig. 4.**
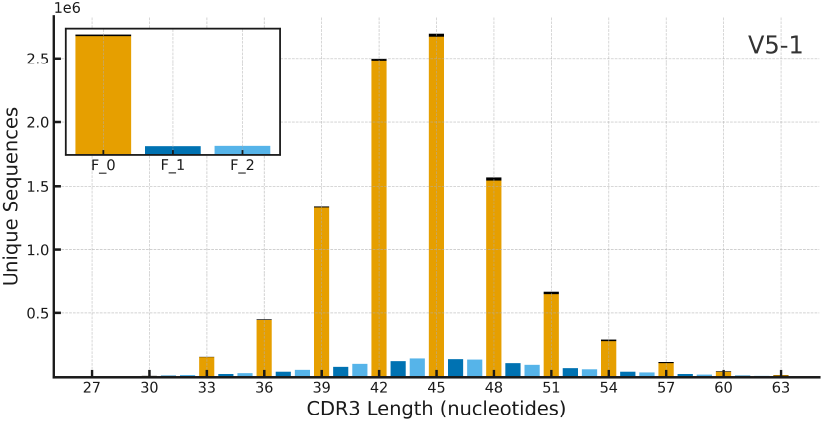
Counts of unique (functional gene) TRBV5-1 sequences by CDR3 length for cohort C1.

**Fig. 5.**
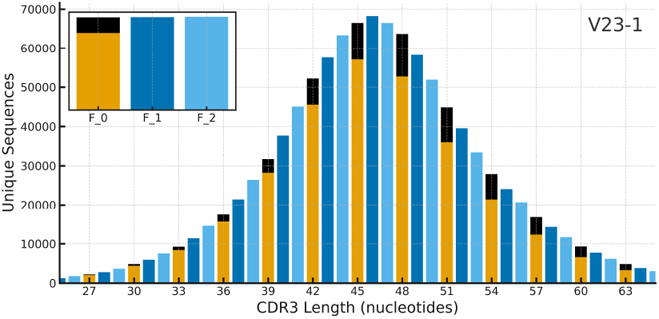
Counts of unique (pseudogene) TRBV23-1 sequences by CDR3 Length for cohort C1.

### Detecting repertoires with unusual TRBV23-1 populations

For pseudogene sequences, the V(D)J model predicts that the ratio of in-frame to out-of-frame sequences should approximate 1:2, and the aggregation over cohort C1 supports this prediction (Figure 5). To evaluate potential deviations from this model, a null hypothesis for repertoire composition was defined. *F*_0_ is partitioned into *F*_0NT_ and *F*_0T_, since a termination codon in CDR3 fundamentally alters the biological consequences of the rearrangement. *F*_0NT_ could in principle yield functional protein, whereas *F*_0T_ is unequivocally non-productive. Including *F*_1_ and *F*_2_, the partitioning yields four compartments for analysis.

Under the null hypothesis, counts across the four compartments within a repertoire were assumed to follow a multinomial distribution with probabilities estimated from the mean compartment proportions across cohorts C1–C9. For each repertoire, a chi-squared test was applied to compare observed and expected counts, yielding a *p*-value (truncated at 10^−20^ for visualization; Figure 6). Repertoires with *p <* 10^−10^ were designated as outliers. The cutoff was chosen pragmatically and appears reasonably well separated from the central mass of the distribution.

**Fig. 6.**
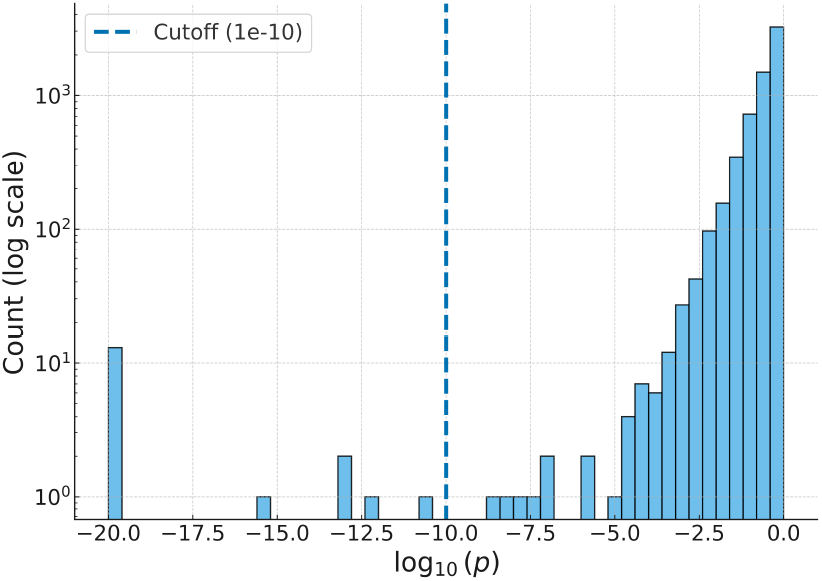
Distribution of log_10_(*p*) values from chi-squared tests across repertoires. Most repertoires clustered within the central mass of the distribution, while a subset showed markedly smaller *p*-values. A pragmatic cutoff of *p <* 10^−10^ was used to designate these as outliers.

Eighteen repertoires met this threshold and are listed in Table 2, with frameshift counts and chi-squared statistics enumerated in Table 3. Of these repertoires, fourteen (D1–D14) form the primary set of outliers considered in the present analysis. The remaining four (S1–S4) are also included in the table for completeness; the rationale for not incorporating them into the main set is deferred to the Discussion.

**Table 2.** Identifiers of outlier repertoires across cohorts.

| Label | Cohort | Sample ID |
| --- | --- | --- |
| D1 | C2 | 03855000014159 |
| D2 | C2 | 03855000013921 |
| D3 | C2 | 03855000012065 |
| D4 | C2 | 03855000011719 |
| D5 | C2 | 03855000012422 |
| D6 | C3 | INCOV012* |
| D7 | C3 | 860011140 |
| D8 | C4 | 435 |
| D9 | C4 | 476 |
| D10 | C5 | MESO121A/B |
| D11 | C6 | R03* |
| D12 | C7 | Pt-09* |
| D13 | C8 | AR_012* |
| D14 | C9 | subj159_D0 |
| S1 | C2 | 03855000014191 |
| S2 | C2 | 03855000014519 |
| S3 | C2 | 03855000013545 |
| S4 | C4 | 392 |

**Table 3.**
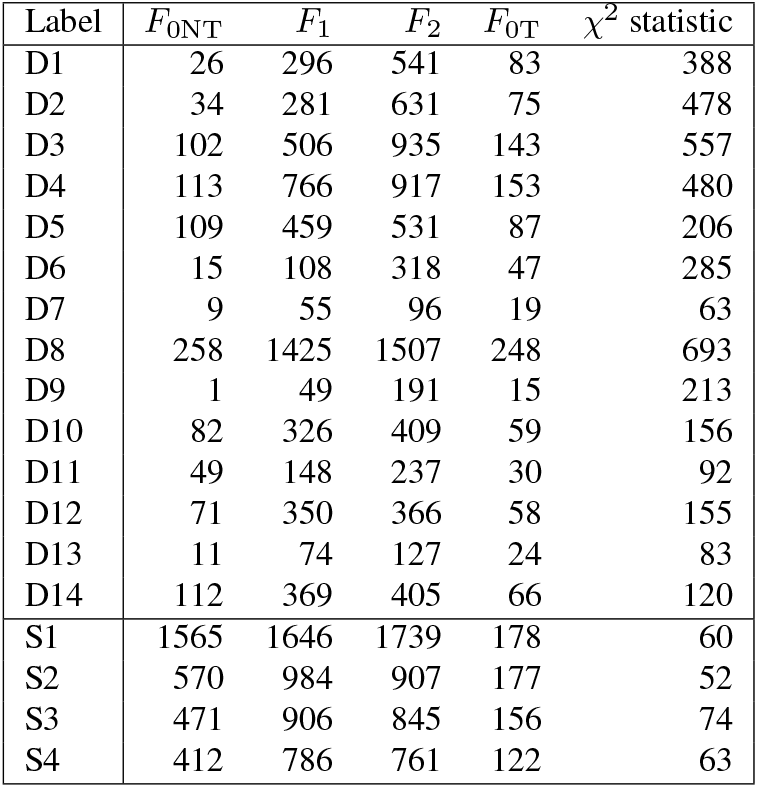
Observed counts in each frame category and chi-square statistics for outlier repertoires.

Combined, cohorts C1–C9 comprise over 6,000 unique individuals. More than 50 additional cohorts were examined but excluded from the present analysis because they contained few individuals, had shallow sequencing depth, or yielded no detectable outliers. These excluded cohorts contributed over 2,000 additional repertoires with no further outliers detected, and estimates of outlier prevalence in this study should be interpreted with this in mind.

Outliers were investigated using the previously described CDR3 length histograms. Figure 7 shows the distribution for repertoire D8. *F*_0NT_ is markedly under-represented relative to *F*_0T_, *F*_1_ and *F*_2_. In contrast, Figure 8 reveals underrepresentation of *F*_0NT_ and (to a lesser degree) *F*_1_ relative to *F*_0T_ and *F*_2_. Histograms for all outliers are available in the Supplementary Data section (Figure S3 and Figure S4).

**Fig. 7.**
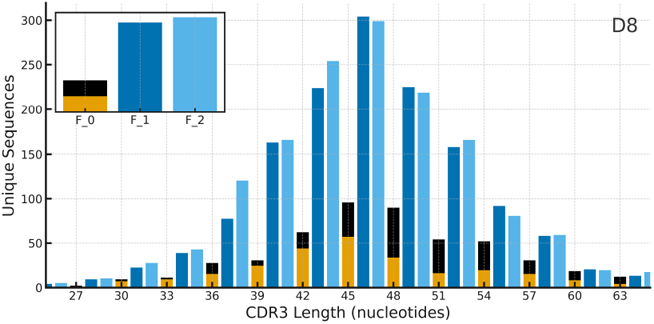
Unique TRBV23-1 sequence counts by CDR3 length for repertoire D8 showing under-representation of *F*_0NT_.

**Fig. 8.**
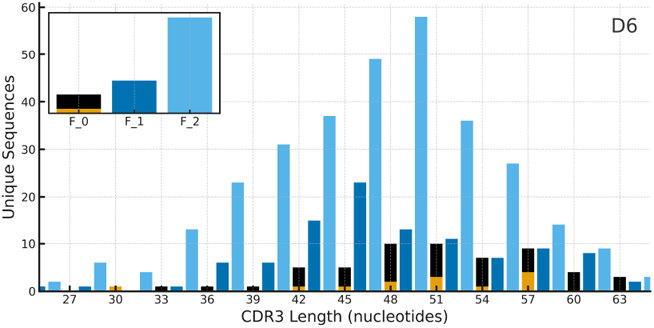
Unique TRBV23-1 sequence counts by CDR3 length for repertoire D6. Both *F*_0NT_ and *F*_1_ are under-represented.

The patterns observed in the outliers motivate plotting *F*_0NT_*/F*_2_ against *F*_1_*/F*_2_ (Figure 9) for all the repertoires in cohorts C1-C9. Lowly sampled repertoires with fewer than 25,000 distinct sequences are excluded from this plot, since TRBV23-1 typically accounts for less than 1% of sequences. At such low coverage, these repertoires would be dominated by sampling noise and cannot generate a meaningful statistical signal.

**Fig. 9.**
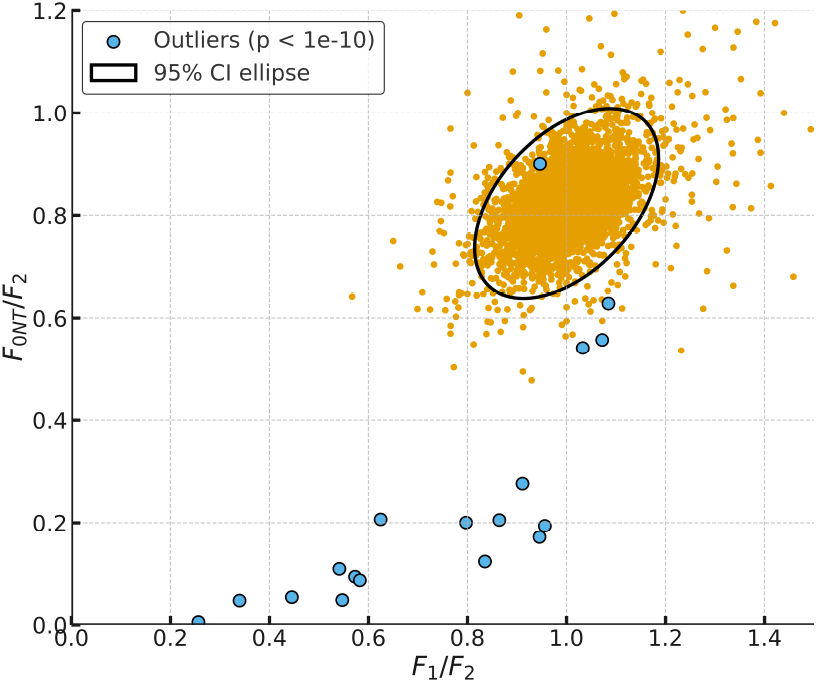
Scatterplot of *F*_0NT_*/F*_2_ versus *F*_1_*/F*_2_ for repertoires from cohorts C1–C9.

Most repertoires cluster into an elliptical cloud oriented along a primary axis *y ≈* 0.8–0.85*x*, rather than *y* = *x*, because *y* is computed from *F*_0*NT*_ rather than *F*_0_ (details in Methods section). A distinct band of outliers, within which a hierarchy is apparent, falls well outside this region. Repertoires on the right of the band exhibit under-representation of *F*_0NT_ relative to *F*_2_ while *F*_1_ remains relatively unaffected. Moving left along the band, under-representation of *F*_0NT_ becomes more pronounced and *F*_1_ shows increasing under-representation as well.

Figure 10 shows an enlarged view of the outlier band with individual repertoires labelled D1–D14, using the labels introduced in Table 2.

**Fig. 10.**
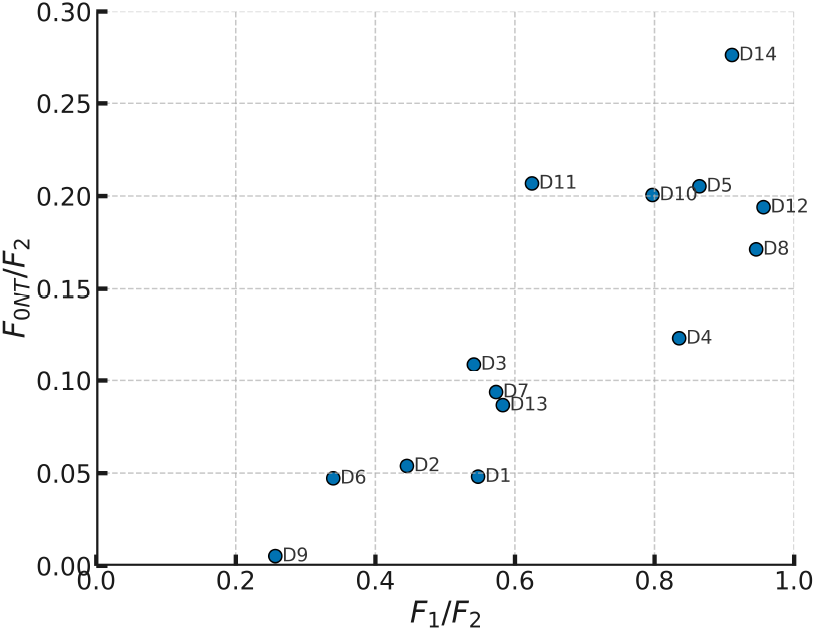
Enlarged view of the frame-ratio scatterplot in Figure 9, showing the band of 14 outlier repertoires (labelled D1–D14, as defined in Table 2).

### Evidence for depletion through *F*_2_ analysis

So far, in all descriptions of frameshift sequence counts, the term *under-representation* has been used to reflect uncertainty over whether the affected frameshift is being depleted or its complementary set enriched. This question cannot be answered with the evidence available so far.

A way to distinguish between these possibilities is provided by the ratio of *F*_2_ sequences assigned to TRBV23-1 relative to all *F*_2_ sequences in a repertoire. For outlier repertoires, this ratio can then be compared with the cohort-wide distribution of the same measure. If the outlier values fall in the lower tail, this would indicate depletion, whereas values in the upper tail would indicate enrichment.

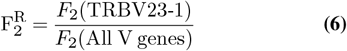

C2, the largest cohort and the one with the greatest number of outliers, was chosen for the initial analysis. Its five outliers (D1–D5) were located on the resulting distribution (Figure 11). Notably, the three repertoires D1–D3, which showed the strongest under-representation in *F*_0NT_ and *F*_1_, fall at CDF values of 0.27, 0.00, and 0.00, respectively—strong evidence that *F*_2_ (and by extension all frames) are being depleted.

**Fig. 11.**
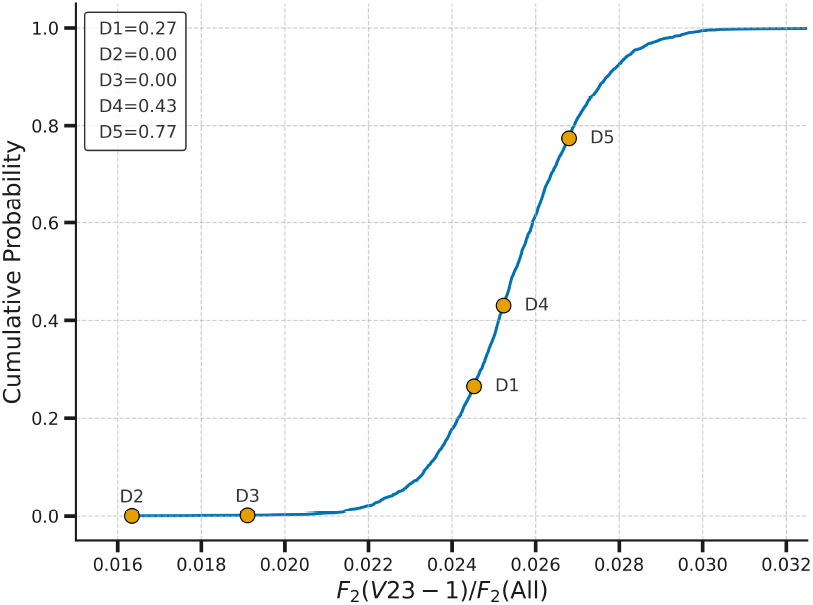
CDF of 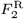 for Cohort C2, with outliers D1–D5 highlighted.

Because experimental conditions such as primer concentration may vary between studies, other outliers must be evaluated within the context of their own cohorts rather than projected onto the C2 distribution. The results for outliers D6, D7, D9 and D13 are most significant, as they fall on the left of the outlier band where both *F*_0NT_ and *F*_1_ are under-represented relative to *F*_2_. If the under-representations truly reflect depletion, these are precisely the cases in which *F*_2_ would also be expected to show depletion. Consistent with this expectation, all four samples fall in the lower tail of their cohort-specific distributions (Table 4; Supplementary Figure S5).

**Table 4.** CDF values for selected outliers, used to test whether TRBV23-1 sequences in *F*_2_ show signs of depletion relative to cohort backgrounds.

| Outlier | CDF |
| --- | --- |
| D6 | 0.00 |
| D7 | 0.03 |
| D9 | 0.00 |
| D13 | 0.08 |

Together, these analyses indicate that the deviations from the null hypothesis observed in TRBV23-1 are best explained by genuine depletion, affecting not only *F*_0NT_ and *F*_1_ but also, in many cases, *F*_2_ (and *F*_0T_).

### RNA-Seq Analysis of TRBV23-1

All analyses so far have been based on DNA sequencing, which captures the full landscape of V(D)J gene usage within PBMC-derived TCR repertoires. To test whether the frame-dependent patterns observed at the DNA level are also reflected at the level of transcript abundance, RNA-seq data from Cohort C10 (Mikelov et al. (15)) were analysed. This cohort provides high-sensitivity measurements of TRBV gene expression. To optimise processing, FASTA files were first filtered for reads containing a sequence unique to TRBV23-1 prior to annotation with igBLAST (see Methods for details). Frame assignment was performed using the CDR3 length distribution, identically to the approach applied in Cohorts C1–C9.

The results, shown in Figure 12, reveal frame-specific RNA levels for TRBV23-1 that are correlated with the depletion hierarchy identified in DNA data. This observation supports an interpretation that post-transcriptional mechanisms contribute to the observed repertoire distortions. The broader implications of this RNA–DNA link are developed further in the Discussion. Future work should extend transcriptomic validation across additional cohorts to test whether the RNA–DNA correspondence observed here is generalizable.

**Fig. 12.**
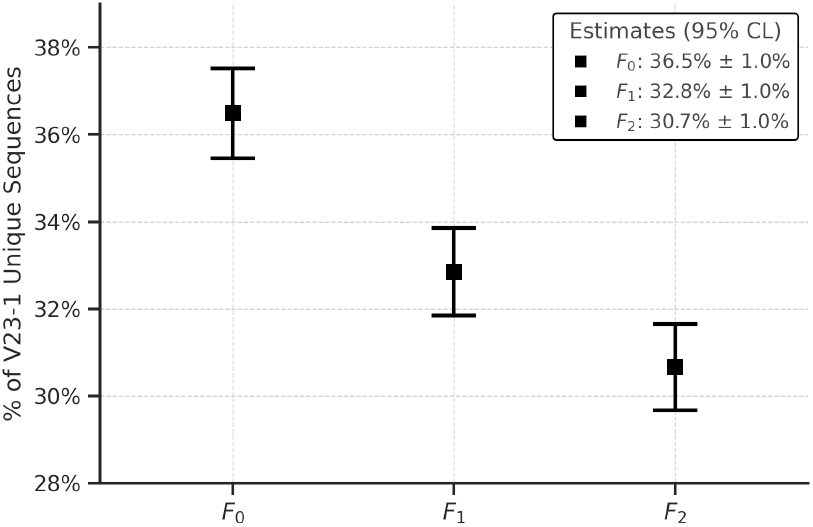
Frame-specific RNA abundances for TRBV23-1 in Cohort C10 (Mikelov et al. (15)). The expression hierarchy mirrors the DNA-based depletion pattern in reverse. This RNA–DNA correspondence provides a key link developed further in the Discussion.

### J-gene family effects on depletion

Having established frame-dependent patterns at both the DNA and RNA levels, J-gene family choice was examined as another possible axis of variation. Each V(D)J rearrangement joins to a J gene from either the J1 or J2 family. Initial inspection suggested that TRBV23-1 sequences joined to J2 genes may experience stronger depletion than those joined to J1 genes.

Direct comparison of J2 usage across individuals is complicated by the fact that the J1/J2 balance varies naturally between repertoires. To normalize for this variation, two quantities were defined for each repertoire: *x*, the overall J2 fraction across all CDR3 sequences for *F*_1,2_, and *y*, the J2 fraction specifically within TRBV23-1 sequences for *F*_all_. A repertoire in which TRBV23-1 follows the global J-family balance will fall along the approximately linear relationship between *x* and *y*, while systematic depletion of J2 within TRBV23-1 will shift points below that line. Figure 13 shows this scatterplot for cohort C2, with outliers from all cohorts superimposed.

**Fig. 13.**
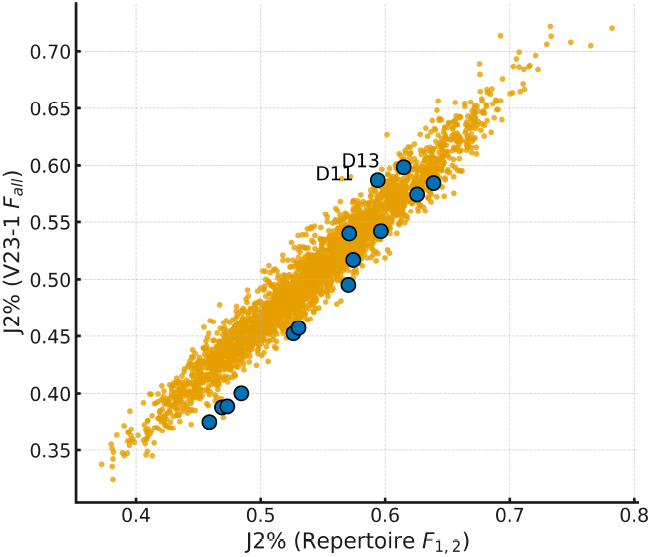
J2 usage in cohort C2 with outliers from all cohorts overlaid. The tendency of outliers to fall along the lower boundary of the cohort distribution indicates selective depletion of J2-associated TRBV23-1 sequences.

Most outliers fall along the lower boundary of the cohort distribution, indicating that TRBV23-1 sequences joined to J2 are disproportionately depleted relative to baseline repertoire usage. In 11 of the 14 outliers this pattern is clear, with D11 and D13 being the main exceptions.

### Other Pseudogenes

Given the clear depletion signal in TRBV23-1, other pseudogenes were also examined for comparable patterns. Among these, TRBV5-3 showed a single clear outlier across cohorts C1–C9, corresponding to outlier repertoire D3 from the TRBV23-1 analysis (Figure 14). RNA data from cohort C10 were used to compute expression levels across frames (Figure 15), and the DNA depletion frameshift pattern in this outlier again matched the RNA abundance pattern. For many pseudogenes, however, DNA counts in the available cohorts were insufficient to reliably identify statistical outliers.

**Fig. 14.**
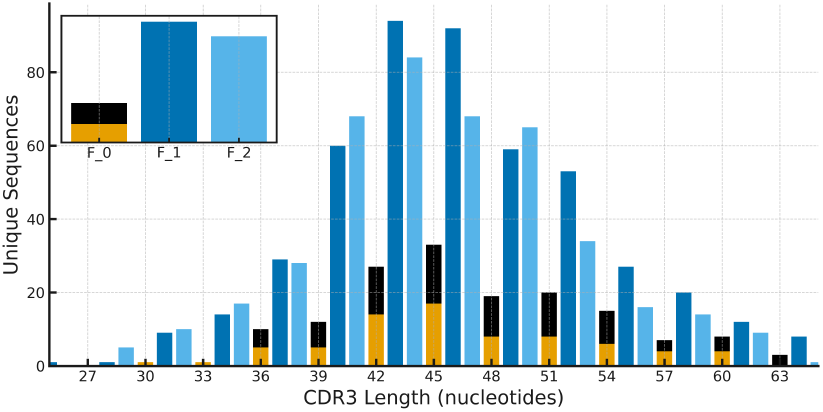
CDR3 length distributions for D3 for TRBV5-3 showing depletion. *F*_0_ is split into *F*_0T_ (top) and *F*_0NT_ (bottom). *F*_0NT_ is once again under-represented.

**Fig. 15.**
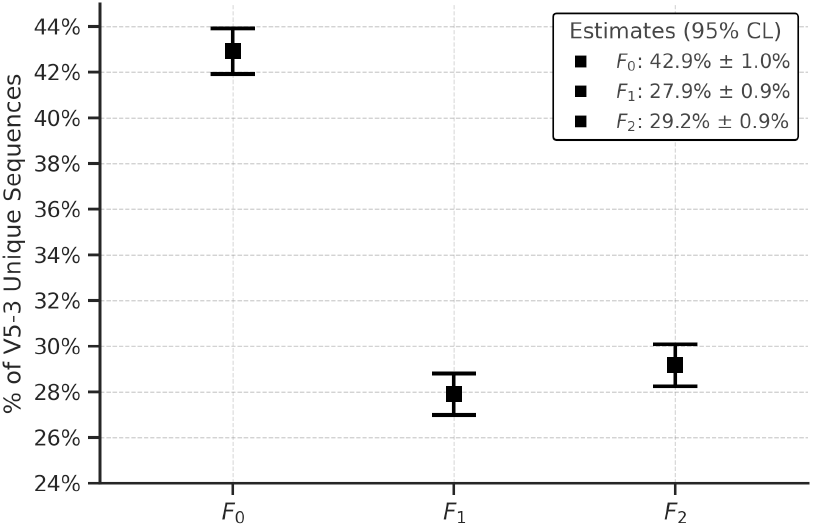
Relative RNA abundances of *F*_0_,*F*_1_ and *F*_2_ for TRBV5-3 in Cohort C10 (Mikelov et al. (15)).

## Discussion

The central finding of this study is that TRBV23-1, a pseudogene assumed to be biologically inert, exhibits frame-dependent depletion in 14 repertoires drawn from 9 cohorts (Table 2). Several possible mechanisms could, in principle, account for the depletion patterns, and the following discussion evaluates these alternatives in turn.

The generation of CDR3 sequences during V(D)J recombination involves multiple independent stochastic steps of nucleotide trimming and insertion (16, 17). By the central limit theorem, repeated convolution of independent random variables drives the distribution of CDR3 lengths toward a normal (Gaussian) distribution. In Fourier space, any higher-frequency periodic structure (such as a frameshift pattern in one random variable) is exponentially suppressed with each convolution, leaving only the universal quadratic term near zero and hence a bell-shaped curve. Indeed, the TRBV23-1 length distribution in cohort C1 (Figure 5) is well approximated by a normal distribution. Frame-dependent generative bias during the recombination process can therefore be ruled out as the cause of the observed depletion patterns.

TRBV23-1 is an open reading frame (ORF) gene with a donor-splice-site mutation (18, 19). TRBV23-1 TCRs have been detected on human T cells through immunostaining (20, 21), suggesting that receptors do reach the cell surface. The earlier in-frame/out-of-frame ratio analysis shows empirically that both types of TRBV23-1 rearrangements are equally likely to be accompanied by a second productive rearrangement. Consequently, the TRBV23-1 TCR transcripts must be expressed at low abundance, insufficient to trigger pre-TCR signalling (22). A possible source of the productive TRBV23-1 transcripts is cryptic or alternative splicing (23–25).

The effect of cryptic or alternative splicing depends on the combined contribution of the reading frame established during V(D)J recombination and the translational phase introduced by the splice donor site. Each splice site, depending on its position relative to the canonical site, would shift the reading frame by 0, 1 or 2 nucleotides. Some sites preserve or restore the global frame, whereas others disrupt it. A single TRBV23-1–bearing T cell can generate a mixture of isoforms, with the different frames *F*_0_, *F*_1_ and *F*_2_ relying on different splice sites to restore register with the J and constant exons.

At first glance, such splicing might offer a plausible explanation for the frameshift depletion patterns in TRBV23-1. In practice, however, any rescue of *F*_1_ and *F*_2_ frames would rewrite conserved regions of the V exon, yielding *β*-chains incompatible with the pre-TCR complex and therefore subject to degradation before reaching the cell surface. More decisively, the hypothesis is contradicted by the observation that even *F*_0T_ sequences, completely beyond the reach of splicing rescue and incapable of producing a TCR under any circumstances, are depleted in the outlier repertoires. In D6, for example, *F*_2_ is depleted by more than 80% while the *F*_0T_*/F*_2_ ratio remains unchanged; in D9, the same ratio is halved.

Antibody recognition of surface expressed receptors can be considered as a source of depletion for *F*_0NT_, but seems unlikely. Productive antibody responses require the assistance of CD4^+^ T cells and HLA-II presentation in professional antigen-presenting cells, which *β* chains are not expected to provide. Extra-follicular responses are possible but are generally weak and transient. Yet again this mechanism cannot explain out-of-frame depletion.

A possible explanation for the depleted outlier band is that a rare TRBV23-1 allele produces a surface-expressed TCR that undergoes negative thymic selection. Accurate interpretation of TCR repertoires depends on reference gene annotations curated by IMGT (18), yet recent studies continue to uncover alleles absent from the database (19). Such a rare allele could therefore remain undocumented. Population-level evidence is incompatible with an allelic explanation, even for the depletion seen in *F*_0NT_. Repertoires D1–D14 show depletions of more than 50% in this compartment, which, if due to a germline variant, would require homozygosity. Among the 6,153 repertoires analyzed, the 14 depleted cases would correspond to a frequency for this rare allele of 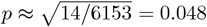. Under Hardy–Weinberg equilibrium, 560 heterozygotes would be predicted, and these should appear as a distinct cluster of partially depleted repertoires around *y≈* 0.4–0.5 in Figure 9. No cluster of this magnitude is observed, and the depleted cases are scattered rather than forming the tight grouping expected for homozygotes. The two alleles of TRBJ2-7 (both productive but of different functionality) provide a real-world example of Hardy–Weinberg equilibrium in T cells (Figure S1 and Figure S2 in the Supplementary data section).

Outliers S2–S4 in Table 2, not discussed until now, could potentially be caused by a rare thymically selected-against allele or alleles since they show less than 50% depletion localized to *F*_0NT_ and could therefore represent the heterozygous case. Equally, these outliers may represent a less advanced stage of the same depletion pattern observed in D1–D14. At present it is not possible to disambiguate between these two possibilities, and for this reason S2–S4 were excluded from the primary analysis. Even if the explanation for these three outliers is allelic, their quantity falls far short of the 560 heterozygotes required by the analysis in the previous paragraph. Outlier S1 is excluded for a different reason. This repertoire shows an unusually low value of *F*_0*T*_ in an individual aged 0–10, with the reduction extending beyond TRBV23-1 to other genes, indicating an anomaly not relevant to the present study.

Insertions or deletions within the conserved V region that do not introduce a premature termination codon can shift the functional reading frame by *±*1 nucleotide. In such cases the productive frame is displaced, with *F*_1_ becoming productive for a −1 or +2 nucleotide shift and *F*_2_ becoming productive for a +1 or −2 nucleotide shift. If the TCR generated were to be the target of negative thymic selection, the DNA-level consequence would be a depletion signature in *F*_1_ or *F*_2_. An example consistent with such an insertion is observed in pseudogene TRBV21-1 for an individual in cohort C2 (Figure S6 in Supplementary Data). Once again, the population-level Hardy–Weinberg expectation implies that heterozygotes would show 50% depletion, producing clusters of partially depleted repertoires in *F*_1_ or *F*_2_, rather than the more depleted scattered multi-frame pattern actually observed.

A cumulative-damage model (described in the Methods section) can plausibly explain the depletion pattern in TRBV23-1 if a suitable frame-dependent damage process can be identified. The process must not involve surface expressed TCRs since frameshifts in the constant region disrupt the transmembrane domain required for anchoring to the plasma membrane (26). HLA-I–mediated presentation of TRBV23-1–derived peptides, which applies equally to all frames, is a potential candidate.

Peptides presented by HLA molecules arise not only from stable proteins undergoing normal turnover but also from defective or misfolded proteins that are rapidly degraded (27). Because the immunopeptidome correlates more closely with the transcriptome than with the proteome (28), even non-productive transcripts can contribute to peptide presentation. RNA-seq data from cohort C10 revealed a clear hierarchy among frames (*F*_0_ *> F*_1_ *> F*_2_) that correlates with the DNA depletion pattern (Figure 12). The empirically observed differences in RNA levels are presumably a consequence of variation in decay efficiency, influenced by sequence context, exon–junction positioning, and the effectiveness of NMD (11, 29, 30), alongside the escape of transcriptional shutdown of *F*_0_.

The peptide-presentation mechanism naturally connects to the cumulative-damage model, since repeated presentation events can drive multi-hit additive cytotoxicity that links transcript abundance to progressive sequence loss. Evidence from modelling and experiments shows that apoptosis due to cytotoxic T cell (CTL) activity often requires the accumulation of multiple sublethal hits rather than a single lethal event (31–34). Such conditions typically arise when CTL–HLA interactions are weak, producing short-lived contacts and only partial responses, further modulated by tissue stiffness and matrix composition (35, 36).

Interacting CTLs release small amounts of perforins and cytokines such as IFN-*γ*, inflicting sublethal damage and amplifying HLA-I presentation, since IFN-*γ* upregulates HLA-I expression (37–39). The peptide-presentation mechanism could, in principle, amplify peptide display and CTL engagement, though the mechanistic details remain unresolved.

A refinement of the cumulative-damage model is that the process may be highly localized in time. CD8 effector responses follow a sigmoidal relationship with the abundance of presented peptide (40–43). TRBV23-1–derived peptides may place cells within the steep portion of this curve, yet without a triggering event, no damage accumulates. If, however, during an immune response a CTL specific for a pathogenic antigen also cross-recognized a TRBV23-1–derived peptide, the coincidence could initiate targeted killing. In this way, a single rare coincidence event can leave a lasting imprint on the repertoire, even though the underlying mechanism remains cumulative damage.

The results from the J-gene family analysis suggest that frameshift depletion does not act uniformly within frame categories but is modulated by J-gene family usage. The data suggest a refinement to the cumulative-damage model whereby the various frames are subdivided into J1 and J2 sub-compartments with different decay rates. J2 damage rates would be higher than those for J1, to capture the asymmetry observed in the data.

In the present work, possible mechanistic explanations for the depletion pattern are explored. HLA-driven peptide presentation is one candidate that aligns with the data, but this remains speculative. Other processes, including small RNA–mediated regulation, affecting T cell behaviour (44, 45), cannot be excluded. The intention of this discussion is not to establish a definitive mechanism, but to provide a robust statistical signal and enumerate testable hypotheses for future work.

Although depletion is only apparent once cells are lost, it is likely that functional impairment occurs earlier, with cumulative-damage compromising T cell performance before overt cell death. TRBV23-1 offered a rare opportunity to observe depletion, as its three frames showed distinct DNA patterns that could be cross-validated with RNA data. Some pseudogenes may display flat RNA profiles, leading to uniform depletion across frames that is difficult to detect, while others may retain an RNA signature but require a higher threshold before depletion is apparent (TRBV5-3 with just a single outlier is a candidate for a higher threshold Figure 14).

The findings underscore a broader point: pseudogenes, long dismissed as inert, may conceal biologically significant patterns. While the mechanistic basis remains unresolved, the results provide strong statistical evidence of non-random depletion and highlight several possible explanatory pathways. Confirming or refuting these will require direct experimental studies. At this stage, the statistical signal is best viewed as evidence for a non-random pattern that motivates further exploration of possible immune regulatory mechanisms.

### Speculative implications

If the peptide-presentation mechanism is correct, it raises the possibility of a self-reinforcing feedback loop. In the scenario where an antigen-responsive T cell happens, by chance, to carry a TRBV23-1 rearrangement alongside its productive rearrangement, three processes would converge. First, infection-driven immune activation elevates IFN-*γ* levels (38, 39). Second, IFN-*γ* may reduce NMD efficiency—an effect reported in pancreatic *β* cells (46) but not yet established to the authors’ knowledge in T cells. Third, clonal expansion of the antigen-responsive cell multiplies TRBV23-1–bearing T cells. Together, these effects could amplify peptide presentation and CTL engagement. While intriguing, this remains a speculative scenario, and further work would be needed to establish any connection to pathological cytokine amplification.

## Methods

### Cohort selection and processing

Ten deep sequence AIRR-seq TCR cohorts were collected and analyzed (Table 5). The nine DNA DSs for the TCR locus were acquired from the open source research database portal immuneACCESS™ published by Adaptive Biotechnologies Corporation, and thus all follow the same Adaptive Biotechnologies immunoSEQ protocol; a multiplex PCR-based assay which allows high-throughput (Illumina) sequencing (56–58). The protocols for the RNA-seq are described in the associated study by Mikelov et al. (15).

**Table 5.** Summary of cohorts used in this study.

| Cohort label | Cohort name | Sequencing protocol | Nucleic acid | Access | Citation |
| --- | --- | --- | --- | --- | --- |
| C1 | Cytomegalovirus (CMV) | immuneSEQ™ | DNA | immuneACCESS™ doi:10.21417/B7001Z | Emerson et al. (47) |
| C2 | SARS-CoV-2 | immuneSEQ™ | DNA | immuneACCESS™ doi:10.21417/RMG2022JCI | Gittelman et al. (48) |
| C3 | immuneCODE(SARS-CoV-2) | immuneSEQ™ | DNA | immuneACCESS™ doi:10.21417/ADPT2020COVID | Nolan et al. (49) |
| C4 | Celiac | immuneSEQ™ | DNA | immuneACCESS™ doi:10.21417/RE2024S | Elyanow et al. (50) |
| C5 | Malignant Pleural Mesothelioma | immuneSEQ™ | DNA | immuneACCESS™ doi: 10.21417/AR2025S | Relevant et al. (51) |
| C6 | Kidney transplant rejection | immuneSEQ™ | DNA | immuneACCESS™ doi:10.21417/sanders2024stm | Sanders et al. (52) |
| C7 | Solid Tumor | immuneSEQ™ | DNA | immuneACCESS™ doi:10.21417/CD2023NM | Chand et al. (53) |
| C8 | Kidney transplant rejection | immuneSEQ™ | DNA | immuneACCESS™ doi:10.21417/TKS2022FI | Sidgel et al. (54) |
| C9 | COVID-19 vaccine | immuneSEQ™ | DNA | immuneACCESS™ doi:10.21417/PAS2021STM | Swanson et al. (55) |
| C10 | SARS-CoV-2 | 5' RACE | RNA | BioProject accession: PRJEB70840 | Mikelov et al. (15) |

All TCR sequence data in the cohorts downloaded from immuneACCESS™ come annotated with component genes using software (igBLAST, MiXCR, etc.) at the authors discretion. The RNA-seq cohort downloaded from the sequence read archive (SRA) has been analysed using igBLAST (v1.21.0) during the course of this study.

For computational efficiency, the FASTA files from cohort C10 were preprocessed to extract candidate TRBV23-1 sequences before running igBLAST. The preprocessing step identified reads containing the nucleotide subsequence ACTGTATCTCTGCG, which was selected because it is (i) unique to TRBV23-1 among all TRBV genes and (ii) positioned near the 3’ end of the V gene, ensuring frequent coverage in the sequencing reads. This filtering substantially reduced file size and computational burden by restricting igBLAST analysis to a small subset of sequences, while discarding rearrangements involving other V genes that were not relevant to the present analysis. Careful checks confirmed that this preprocessing step did not introduce any bias in frameshift distribution. After filtering, igBLAST (v1.21.0) was applied, yielding 78,481 total rows, of which 77,735 were annotated as TRBV23-1. Unique sequences were then counted for each repertoire across the three frameshifts and summed over all 134 repertoires, resulting in counts of 3,057, 2,752, and 2,569 for *F*_0_, *F*_1_, and *F*_2_, respectively. For TRBV5-3 a similar procedure was adopted, preprocessing with the nucleotide sequence GCCTTGGAGCTGGGGGAC, resulting in counts of 4,026, 2,617 and 2,736 for *F*_0_, *F*_1_, and *F*_2_, respectively.

All cohort references but C4 refer to their respective studies for which the associated data was collected. The reference to C4 refers to the immuneACCESS™ webpage where it was made available (https://clients.adaptivebiotech.com/pub/elyanow-2024-s subject to Adaptive Biotechnologies publishing policies) with an associated study to be determined.

Column three of Table 6 shows the number of samples used after combining multiple samples from the same individual into the same repertoire to increase statistical precision and avoid outlier duplication. The number of original samples downloaded for each cohort and the final number of repertoires analysed following merger are shown in Table 6. Note that the immuneCODE™ (C3) cohort used is release 002.

**Table 6.** Number of repertoires analysed per cohort, before and after merging same person repertoires where identifiable. Totals for cohorts C1-C9 are also included.

| Cohort | Samples | Merged Samples | Outliers |
| --- | --- | --- | --- |
| C1 | 786 | 786 | 0 |
| C2 | 2889 | 2889 | 5 |
| C3 | 1414 | 1304 | 2 |
| C4 | 520 | 520 | 2 |
| C5 | 40 | 20 | 1 |
| C6 | 631 | 56 | 1 |
| C7 | 30 | 15 | 1 |
| C8 | 327 | 195 | 1 |
| C9 | 363 | 234 | 1 |
| **C1-C9** | **7134** | **6153** | **14** |
| C10 | 134 | 134 | N/A |

### Modelling the expected in-/out-of-frame ratio

The expected ratio of in-frame to out-of-frame rearrangements in the naïve T cell repertoire after thymic selection was modelled, incorporating multiple biological and technical factors. Two key parameters are central to the model: *p*_2_, the probability that a non-productive first rearrangement undergoes a second recombination attempt; and *p*_T_, the probability that an in-frame rearrangement contains a PTC in the CDR3 region.

For each V gene *v*, structural functionality is defined as the probability that an in-frame rearrangement lacking a PTC yields a functional receptor. This quantity is represented by

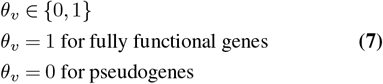

Even for functional genes, in-frame rearrangements may still be non-productive if they contain a PTC in the CDR3, with probability *p*_T,v_. The effective productivity of an in-frame rearrangement of gene *v* is therefore

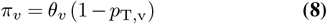

Following recombination, expressed productive chains in *F*_0_ survive thymic selection with probability *σ*_*v*_ ∈ [0, 1]. Thymic selection may also act on the first recombination attempt even when it fails and is subsequently rescued by a second attempt. These survival factors apply specifically to failed first arrangements and may differ by frame. To capture this, frame-specific survival probabilities 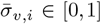 are introduced for each gene *v* and frameshift *i*.

Each gene *v* also has a recombination usage frequency *U*_*v*_, where ∑_*v*_ *U*_*v*_ = 1. Combining this with the effective productivities *π*_*v*_ gives the cohort-average productivity

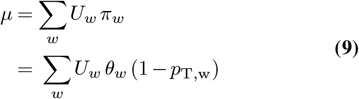

Rearrangements that fail on the first attempt may undergo a second recombination with probability *p*_2_, assumed to be gene-independent. Frame generation probabilities (in-frame = 1*/*3, out-of-frame = 2*/*3) are built into the constants below, recognising that in reality these values may vary slightly depending on the CDR3 length distribution.

The statistic of interest is the gene-specific ratio

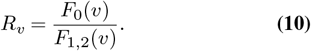

The contributions of in-frame and out-of-frame rearrangements can be expressed as

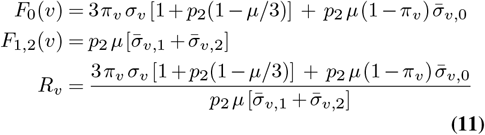

Several special cases illustrate the behaviour of this expression. For pseudogenes (*π*_*v*_ = 0) and

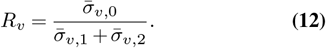

With no frame-dependent filtering, this reduces to *R*_*v*_ = 0.5.

Another special case concerns functional genes assuming a second recombination attempt is guaranteed (*p*_2_ = 1)

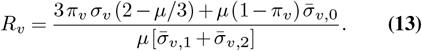

Without making any assumptions about *p*_2_, consider that a fraction *f* of V genes are pseudogenes, all functional genes have the same termination probability *p*_T_, and frame filters are equal 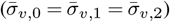. Then

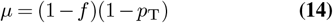

and the ratio for fully functional V genes reduces to

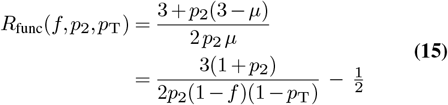

This simplified form provides a natural baseline expectation against which empirical repertoire data can be compared. In particular, the predicted range of *R*_func_ aligns closely with the V-gene–specific ratios observed in cohort C1 (Figure 1), supporting the validity of the model before introducing further refinements such as frame-specific effects.

### Expected geometry of frame ratios under the multi-nomial null

Under the null hypothesis, repertoire counts (*F*_0NT_, *F*_0T_, *F*_1_, *F*_2_) follow a multinomial distribution with probabilities (*q*_0NT_, *q*_0T_, *q*_1_, *q*_2_). For a repertoire of size *N*, this implies

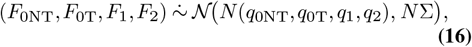

where Σ is the covariance matrix of the multinomial.

Each recombination attempt generates frame classes with equal probability 1*/*3. However, only a fraction (1 − *p*_T_) of *F*_0_ sequences avoid a premature termination codon in the CDR3 and contribute to *F*_0NT_, while the remainder contribute to *F*_0T_.

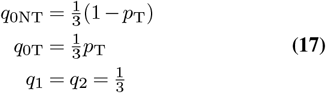

For realistic termination codon rates *p*_T_ *≈* 0.15–0.20, this yields *q*_0NT_ *≈* 0.27–0.28, compared to *q*_1_ = *q*_2_ *≈* 0.33, so that *q*_0NT_ *≈* 0.8–0.85 of *q*_1_ and *q*_2_.

To visualise frame usage, the ratios *x* = *F*_1_*/F*_2_ and *y* = *F*_0NT_*/F*_2_ are used, with *F*_0T_ treated as an additional multinomial category contributing to the covariance structure.

By the delta method, (*x, y*) is asymptotically bivariate normal with mean (*q*_1_*/q*_2_, *q*_0NT_*/q*_2_) and covariance Σ_*R*_*/N* for a fixed 2 *×* 2 matrix Σ_*R*_. Iso-density contours of a bivariate normal are ellipses. With *p*_T_ = 0.18

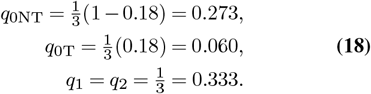

The centre of the iso-density ellipses are therefore located at

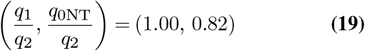

The primary axis of the ellipses should lie on a line *y* = (*q*_0NT_*/q*_1_)*x ≈* 0.8–0.85*x*. The length of the axes of the ellipses shrink like 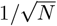, so repertoires with larger numbers of sequences fall into a tighter elliptical cloud.

### A cumulative-damage model explains sequential depletion across frameshift classes

Across the cohorts analysed, no significant bias in the relative abundance of *F*_0_, *F*_1_, and *F*_2_ was observed, consistent with balanced generative and survival rates for these populations. However, as already shown, in a subset of outliers, this balance was disrupted, with some repertoires showing selective depletion in *F*_0_, others in both *F*_0_ and *F*_1_, and the most extreme cases showing loss across all three frameshift classes. To explain this ordered pattern of depletion, the analysis considered whether a simple mathematical model could explain the sequential loss of frameshift populations. In this simplified model, no distinction is made between *F*_0T_ and *F*_0NT_.

T cell survival was modelled as a cumulative-damage process, in which cells sustain random damage over time, and die once a fixed threshold of damage is exceeded. In this framework, the effective damage rate is linked to the frame context, with *F*_0_ cells experiencing the highest rate, followed by *F*_1_ and then *F*_2_. This ordering naturally produces a staggered cascade of loss, with *F*_0_ reaching the threshold first, then *F*_1_, and ultimately *F*_2_.

When viewed across repertoires, this model can explain why most repertoires show no bias in the relative abundance of *F*_0_, *F*_1_, and *F*_2_: in these cases, cumulative damage has not yet advanced far enough to cause measurable depletion in any subset. In contrast, outlier repertoires reveal the sequential signature of the process as some show depletion confined to *F*_0_, others show loss in both *F*_0_ and *F*_1_, and in the most extreme cases in *F*_0_, *F*_1_ and *F*_2_. By linking the effective hit rate to the frameshift context, a simple mechanistic basis for the observed repertoire patterns is obtained.

The attrition of *F*_0_, *F*_1_, *F*_2_ was modelled using an Erlang survival process. The Erlang form,

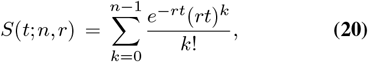

describes the probability that a clone survives after cumulative stress *t*, given that *n* independent “hits” are required for loss and that hits arrive as a Poisson process of rate *r*. This formulation provides a natural way to capture the idea that cellular loss occurs only after repeated events rather than instantaneously. Different frames (*F*_0_, *F*_1_, *F*_2_) were assigned distinct rate parameters *r*_*f*_ such that their half-lives occur at 40, 80, and 120 cumulative units, respectively. These values are purely illustrative and chosen to demonstrate the qualitative separation of attrition dynamics between frames, and should not be interpreted as direct fits to empirical data.

Figure 16 shows the resulting survival curves for the three frames, illustrating their characteristic attrition rates under the cumulative damage model. To visualize survival at discrete checkpoints, Figure 17 summarizes the surviving fractions of *F*_0_, *F*_1_, *F*_2_ at *t* = 0, 40, 80, and 120. Although presented here in a discrete-event form, the Erlang model is easily extended to a continuous attrition framework, where survival decays smoothly with stress intensity or duration.

**Fig. 16.**
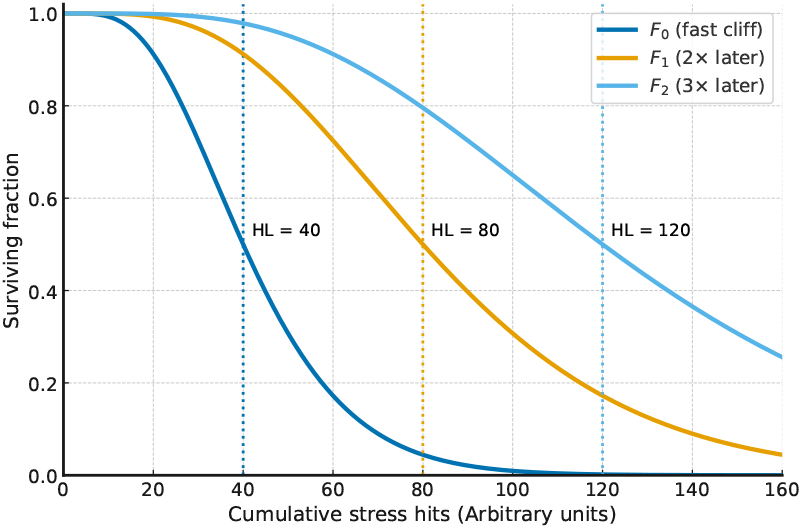
Cumulative damage model showing survival curves for frames *F*_0_, *F*_1_, and *F*_2_ as a function of cumulative stress hits (arbitrary units), each with a distinct attrition rate.

**Fig. 17.**
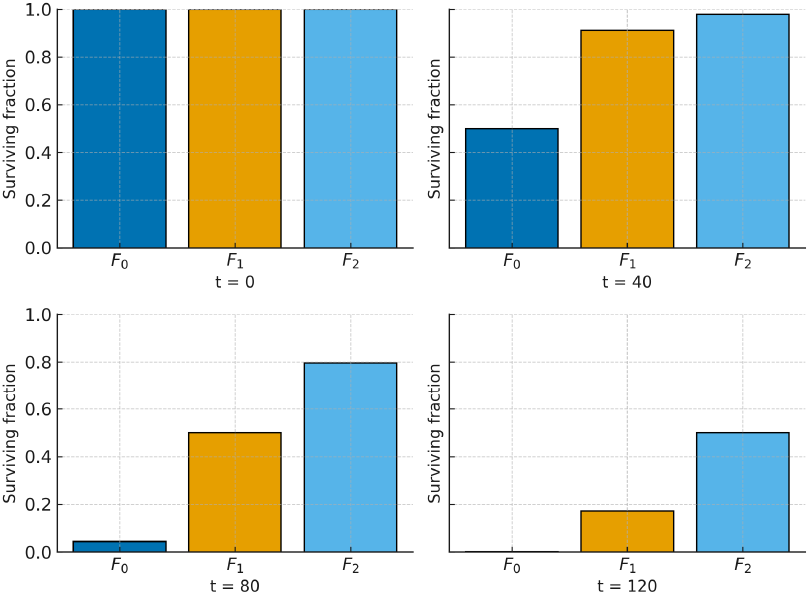
Surviving fractions of frames *F*_0_, *F*_1_, and *F*_2_ at *t* = 0, 40, 80, and 120 (arbitrary units) under a cumulative damage model with different attrition rates for each frame.

## Data Availability

The cohorts used for the analysis, alongside relevant immuneACCESS™ DOIs and BioProject accessions can be found in Table 5. The code for the analyses and the results are available in the GitHub repository (https://github.com/parsimonial/TiggBektashiBrown.git). The IMGT TRBV references used can be found at https://www.imgt.org.

## Declarations

### Ethics approval and consent to participate

Not applicable.

### Consent to publish

Not applicable.

### Author contributions

**JT:** Conceptualisation (project design, framing of research questions, motivating and constructing mathematical models), Methodology, Formal analysis, Writing – original draft, Writing – review and editing.

**ABB:** Conceptualisation (integration of biological context and development of explanatory models), Writing – original draft, Writing – review and editing.

Both authors contributed to the final manuscript and approved the submitted version.

