## Supplemental Data for "Frame-specific depletion of the TRBV23-1 pseudogene in human TCR repertoires: Quantitative evidence and possible biological explanations"

### Supplementary Data

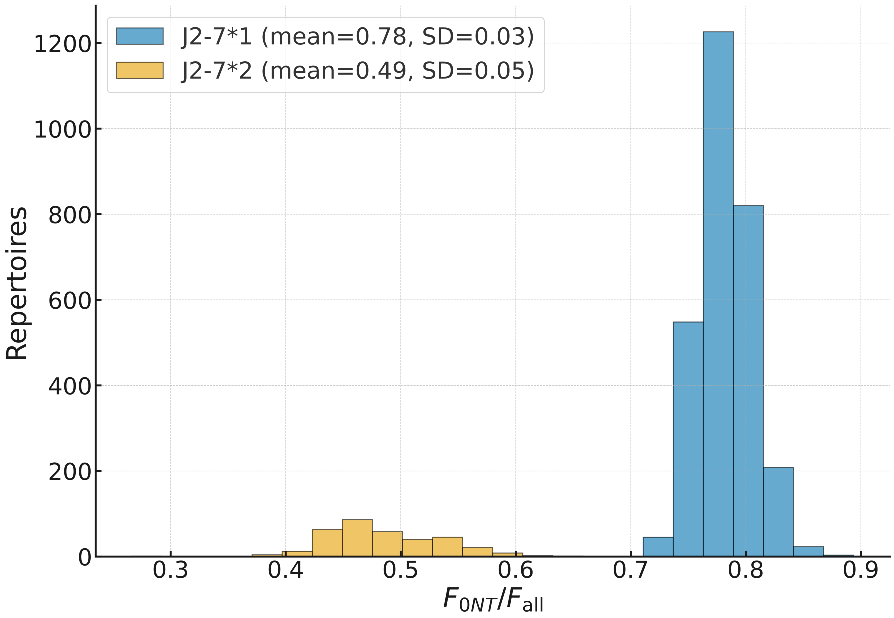


**Fig. S1.** Distribution of $F_{\mathrm{0NT}}/F_{\mathrm{all}}$ in cohort C2 for the two functionally distinct TRBJ2-7 alleles, computed separately for TRBJ2-7*1 and TRBJ2-7*2. Here the alleles are resolved directly from sequencing reads, so their presence need not be inferred from functional patterns.

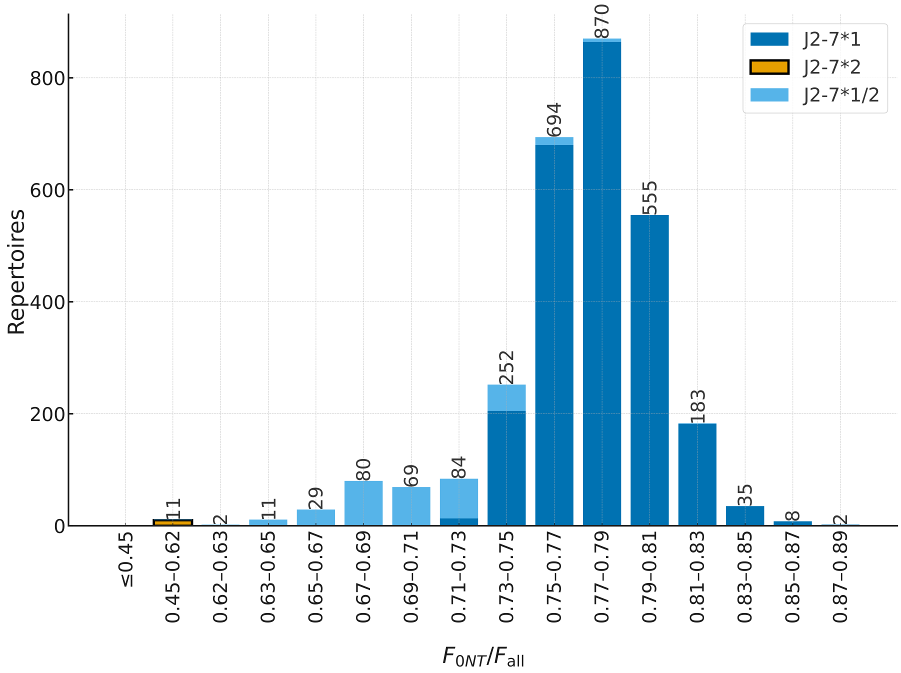


**Fig. S2.** Distribution of $F_{\mathrm{0NT}}/F_{\mathrm{all}}$ for TRBJ2-7 in cohort C2, with the two resolved alleles pooled to mimic a single label, yielding a trinomial distribution whose Hardy–Weinberg weights reveal the presence of two alleles with different functionality.

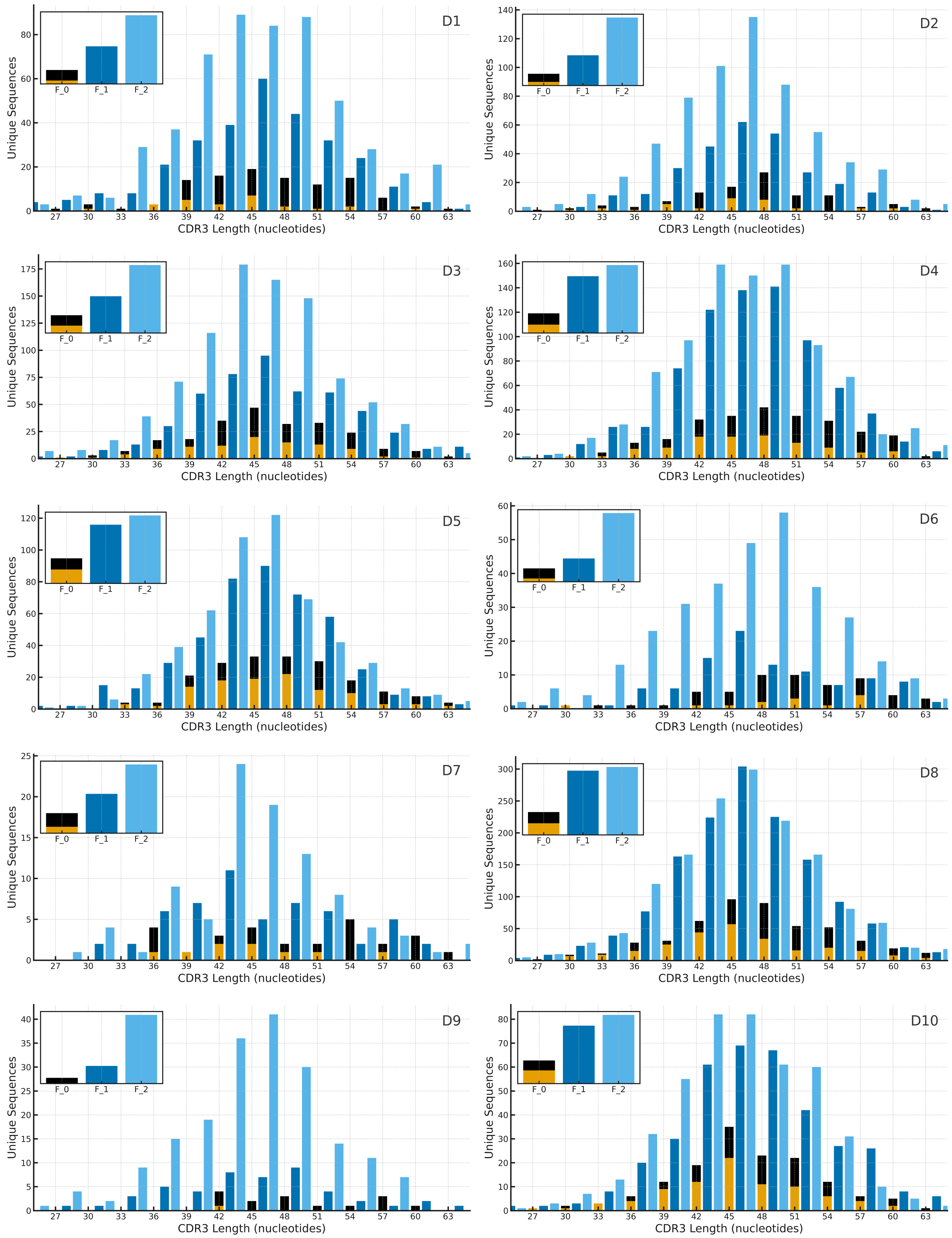
D1
D2
D3
D4
D5
D6
D7
D8
D9
D10
Unique Sequences
CDR3 Length (nucleotides)
F_0
F_1
F_2
27
30
33
36
39
42
45
48
51
54
57
60
63

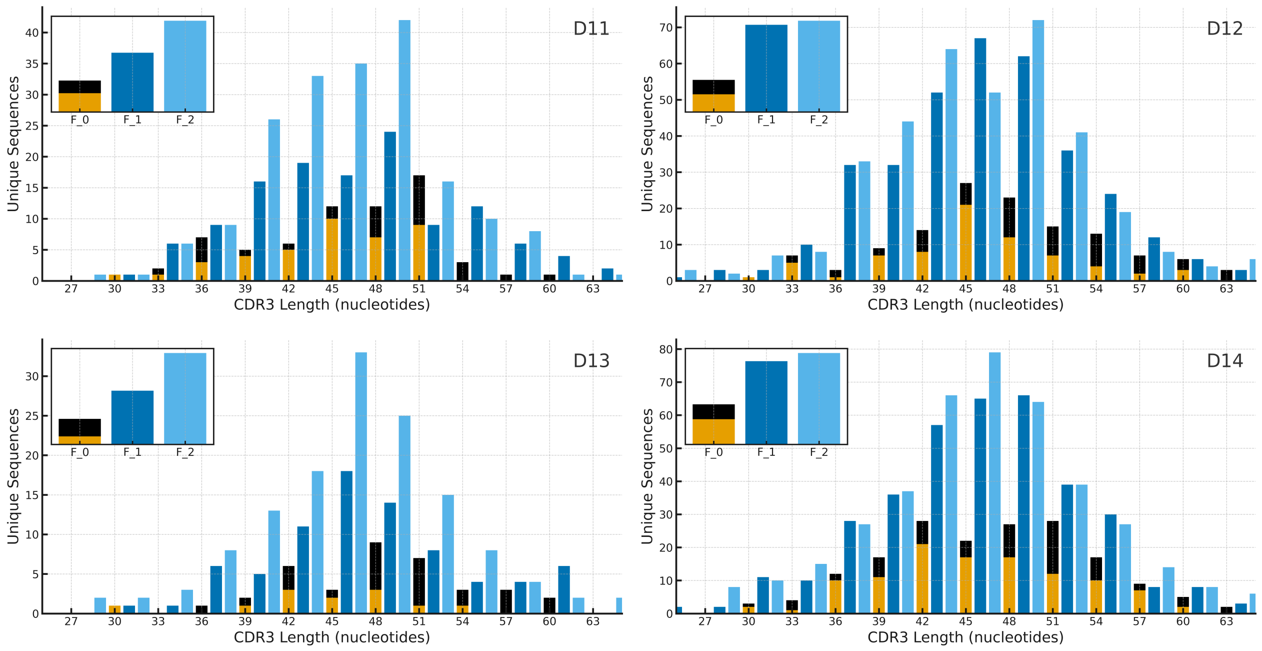


**Fig. S3.** Unique TRBV23-1 sequence counts by CDR3 length for outliers D1–D14. $F_0$ is split into $F_{0\mathrm{T}}$ (top) and $F_{0\mathrm{NT}}$ (bottom).

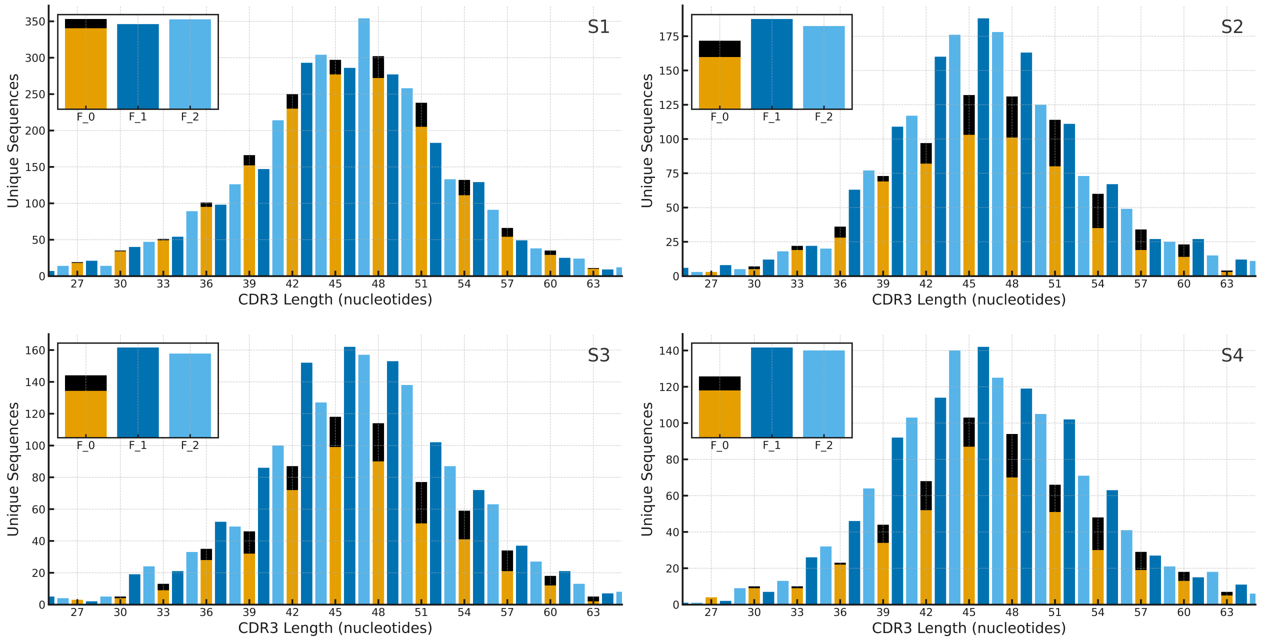


**Fig. S4.** Unique TRBV23-1 sequence counts by CDR3 length for excluded outliers S1–S4. $F_0$ is split into $F_{0\mathrm{T}}$ (top) and $F_{0\mathrm{NT}}$ (bottom). S2-S4 may be depletion or allelic thymic selection. S1 (0–10 year-old individual) shows reduced termination codons across multiple V genes, including TRBV23-1.

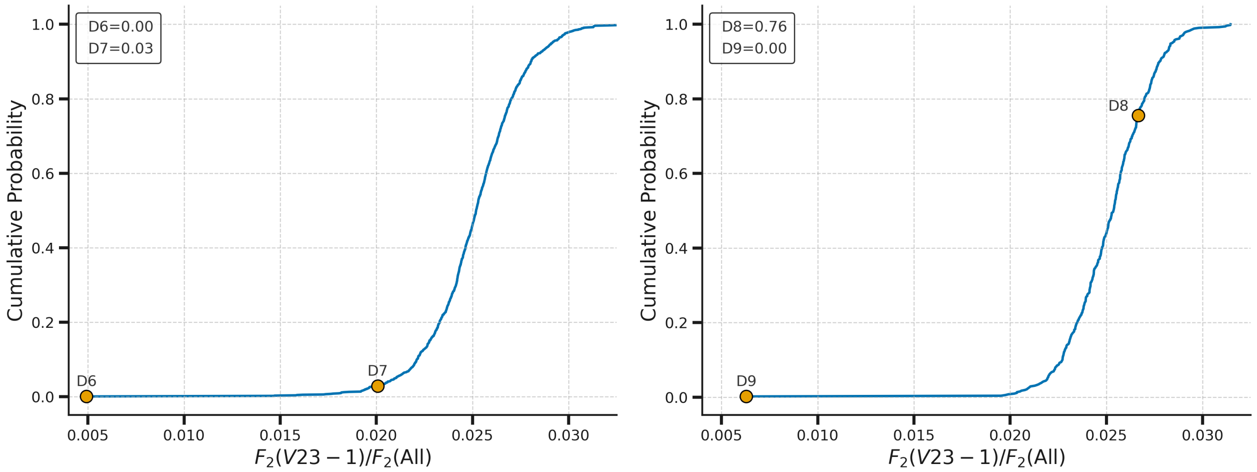


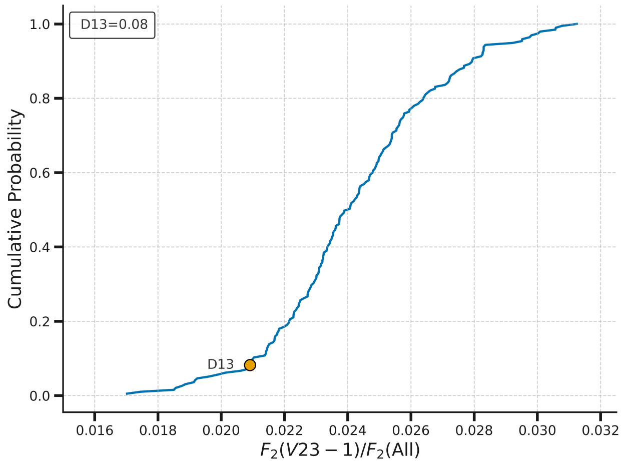


**Fig. S5.** CDF of $F_2^{\mathrm{R}}$ with outliers for Cohorts C3 (D6, D7), C4 (D8,D9) and C8 (D13). D6, D7, D9 and D13 lie at the lower end of their cohort distributions showing depletion of the $F_2$ compartment. Cohort C8 comprises only 195 samples, so its CDF is less smooth.

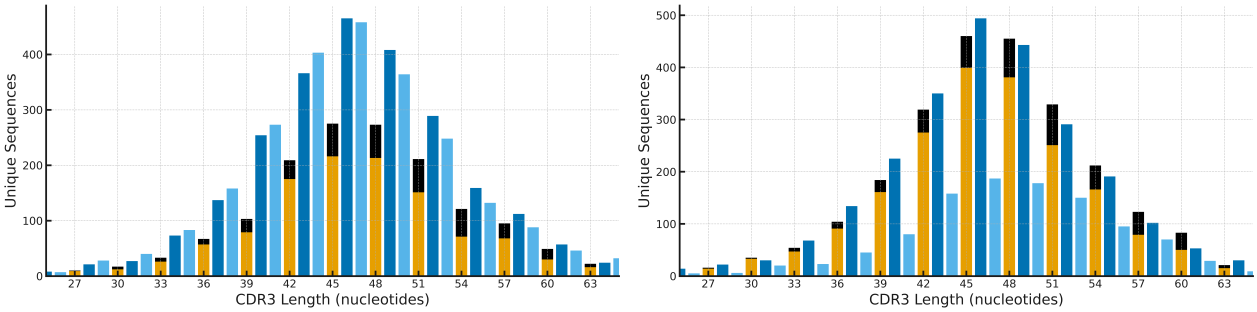


**Fig. S6.** CDR3 length distributions for TRBV21-1 in two samples from Cohort C2. $F_0$ is split into $F_{0\mathrm{T}}$ (top) and $F_{0\mathrm{NT}}$ (bottom). Sample 03855000012466 (left) shows a pattern consistent with a thymically selected against allele causing depletion in $F_0$ Sample 03855000012402 (right) shows a pattern consistent with a thymically selected insertion causing depletion in $F_2$. The sequence window does not extend far enough into the V region to resolve the precise nucleotide changes.
